# Hepatic lipid overload potentiates biliary epithelial cell activation via E2Fs

**DOI:** 10.1101/2022.07.26.501569

**Authors:** Ece Yildiz, Gaby El Alam, Alessia Perino, Antoine Jalil, Pierre-Damien Denechaud, Katharina Huber, Lluis Fajas, Johan Auwerx, Giovanni Sorrentino, Kristina Schoonjans

## Abstract

During severe or chronic hepatic injury, biliary epithelial cells (BECs), also known as cholangiocytes, undergo rapid reprogramming and proliferation, a process known as ductular reaction (DR), and allow liver regeneration by differentiating into both functional cholangiocytes and hepatocytes. While DR is a hallmark of chronic liver diseases, including advanced stages of non-alcoholic fatty liver disease (NAFLD), the early events underlying BEC activation are largely unknown. Here, we demonstrate that BECs readily accumulate lipids upon fatty acid (FA) treatment in BEC-derived organoids, and during high-fat diet feeding in mice. Lipid overload induces a metabolic rewiring to support the conversion of adult cholangiocytes into active BECs. Mechanistically, we found that lipid overload unleashes the activation of the E2F transcription factors in BECs, which drives cell cycle progression while promoting glycolytic metabolism. These findings demonstrate that fat overload is sufficient to initiate a DR, without epithelial damage, and provide new insights into the mechanistic basis of BEC activation, revealing unexpected connections between lipid metabolism, stemness, and regeneration.

## Introduction

Under physiological conditions, the hepatic epithelium, composed of hepatocytes and BECs (or cholangiocytes), is non-proliferative. Yet upon injury, these two cell types are capable of rapidly changing their phenotype from quiescent to proliferative, contributing to the prompt restoration of damaged tissue (Gadd et al., 2020; Michalopoulos, 2014; Miyajima et al., 2014; Yanger and Stanger, 2011). However, in chronic liver injury, characterized by impaired hepatocyte replication, BECs weigh-in and serve as the cell source for regenerative cellular expansion through the DR process (Choi et al., 2014; Deng et al., 2018; Español–Suñer et al., 2012; Huch et al., 2013; Lu et al., 2015; Raven et al., 2017; Rodrigo-Torres et al., 2014; Russell et al., 2019).

The molecular basis by which BECs expand during the DR has been extensively studied in chemical models of biliary damage and portal fibrosis using the chemical 3,5-diethoxycarbonyl-1,4-dihydrocollidine (DDC). Several signaling pathways involving YAP (Meyer et al., 2020; Pepe-Mooney et al., 2019; Planas-Paz et al., 2019), mTORC1 (Planas-Paz et al., 2019), TET1-mediated hydroxymethylation (Aloia et al., 2019) and NCAM1 (Tsuchiya et al., 2014) have been reported to drive this process. Importantly, DR has also been observed in late-stage NAFLD patients with fibrosis and portal inflammation (Gadd et al., 2014; Sato et al., 2018; Sorrentino et al., 2005). NAFLD, one of the most common chronic diseases, initiates with increased lipid accumulation, a stage called steatosis (Paschos and Paletas, 2009). This pathology progresses into inflammation and fibrosis that can cause cirrhosis and hepatocellular carcinoma, which are the most frequent liver transplantation indications (Byrne and Targher, 2015). YAP has been found to be activated in BECs in fibrotic livers but not in steatosis (Machado et al., 2015), suggesting that YAP activation is necessary to support DR in the late fibrotic NAFLD stages, and thus, leaving the early molecular mechanisms of BEC activation unexplored.

BEC-derived organoids (BEC-organoids) can be established from intrahepatic bile duct progenitors and exhibit a DR-like signature, representing a promising *in vitro* approach to study regenerative mechanisms and therapies (Huch et al., 2015, 2013; Li et al., 2017; Okabe et al., 2009; Shin et al., 2011; Sorrentino et al., 2020). These self-renewing bi-potent organoids are capable of expressing stem cell/progenitor markers and of differentiating into functional cholangiocyte- and hepatocyte-like lineages, which can engraft and repair bile ducts (Hallett et al., 2022; Sampaziotis et al., 2021) and improve liver function when transplanted into a mouse with liver disease (Huch et al., 2015, 2013; Li et al., 2017).

Here, we used BEC-organoids and BECs isolated from chow diet (CD)- or high-fat diet (HFD)-fed mice and reported that they are affected by acute and chronic lipid overload, one of the initial steps of NAFLD. Lipid accumulation turns BECs from quiescent to proliferative cells and promotes their expansion through the E2F transcription factors and the concomitant induction of glycolysis. These observations hence attribute a pivotal role to E2Fs, regulators of cell cycle and metabolism, in inducing BEC activation before the late fibrotic stage of NAFLD.

## Results

### BECs and BEC-organoids efficiently accumulate lipids in vivo and in vitro

To gain insight into how chronic lipid exposure, an inducer of liver steatosis, affects biliary progenitor function *in vitro*, we incubated single BECs with a fatty acid mixture (FA mix) of oleic acid (OA) and palmitic acid (PA) – the two most abundant FAs found in livers of NAFLD patients (Araya et al., 2004), for 7 days and allowed BEC-organoid formation (Figure 1A). Surprisingly, we observed that BEC-organoids efficiently accumulated lipid droplets in a dose-dependent manner (Figure 1B) and this process did not affect organoid viability (Figures 1C-D). To investigate how cells adapt their metabolism to lipid overload, we monitored the expression of several genes involved in lipid metabolism, including *Scd1* (*de novo* lipogenesis) (Figure 1E), *Hmgcs2* (ketogenesis), *Pdk4* (inhibition of pyruvate oxidation), and *Aldh1a1* (prevention against lipid peroxidation products) (Figure 1F) and found it to be affected by FA addition. These results suggest that BEC-organoids actively reprogram their metabolism to cope with aberrant lipid overload.

**Figure 1.**
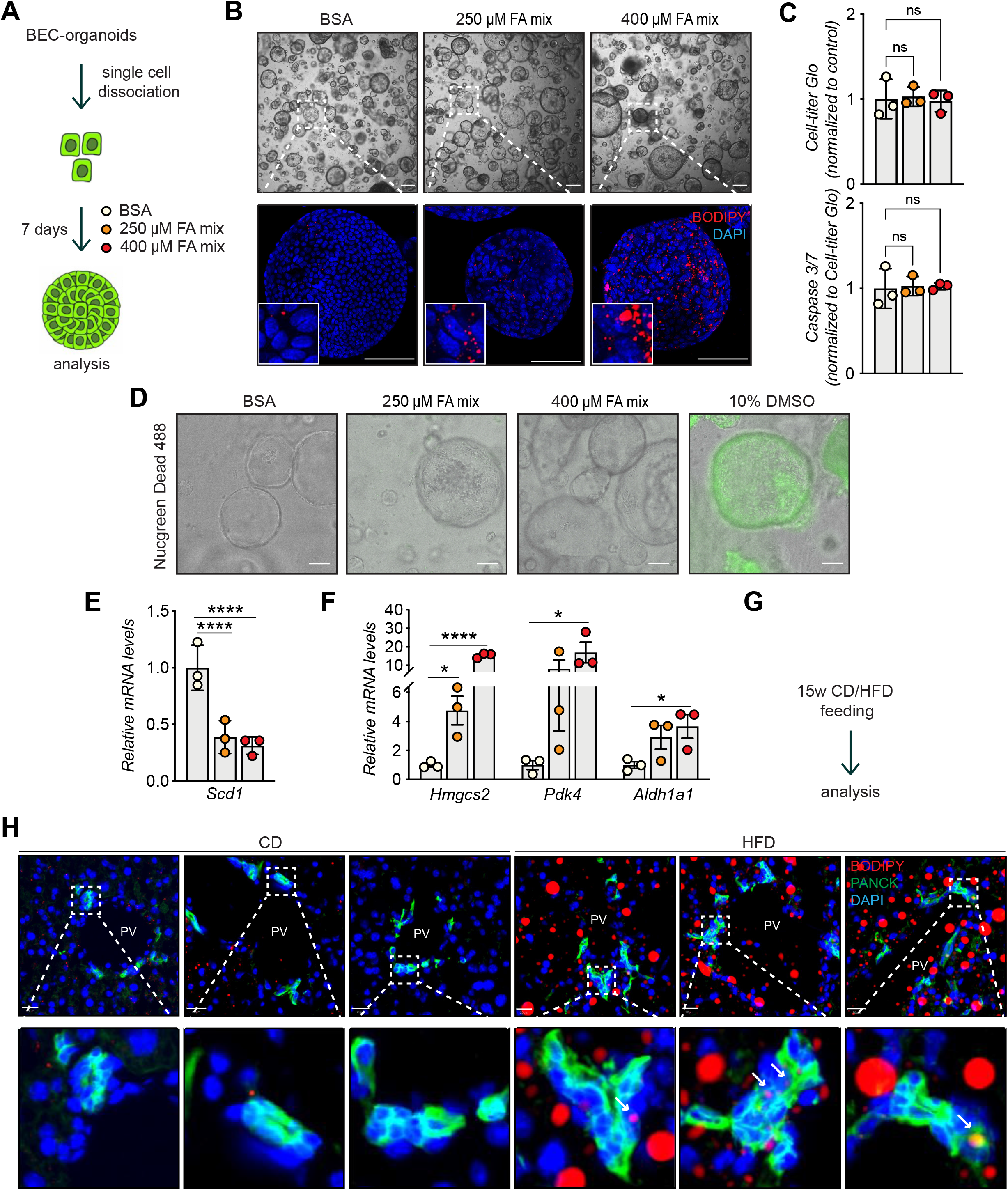
BECs accumulate lipids. **(A)** Schematic depicting fatty acid (FA) treatment of BEC-organoids *in vitro*. **(B)** Representative brightfield and immunofluorescence (IF) images of lipids (BODIPY) in control (BSA) and FA-treated organoids. Close-up IF images were digitally zoomed in four times. n=3. **(C)** Cell-titer Glo and Caspase 3/7 activity measurement for viability and apoptosis detection relative to panel A. n=3. **(D)** Representative Nucgreen Dead 488 staining as composite images from brightfield and fluorescent microscopy. n=3. **(E-F)** Quantification of *Scd1* (**E**) and *Hmgcs2*, *Pdk4,* and *Aldh1a1* (**F**) mRNA in control (BSA) and FA-treated organoids. n=3. **(G)** Schematic depicting CD and HFD feeding *in vivo*. **(H)** Representative images for co-staining of BODIPY and PANCK, relative to panel G. Close-up IF images were digitally zoomed in four times. n=5. Data are shown as mean ± SD. Absence of stars or ns, not significant (p > 0.05); *p < 0.05; ****p < 0.0001; one-way ANOVA with Dunnett’s test (C), and Fisher’s LSD test (E, F) was used. PV, portal vein. Arrowheads mark bile ducts. Scale bars, 200 μm (B-brightfield), 100 μm (B-IF, D) and 20 μm (H-I). The following figure supplements are available for figure 1: Figure supplement 1. Further characterization of lipid accumulation in BECs.

To determine whether the observed phenotype was preserved in fully formed organoids, we treated already established BEC-organoids with the FA mix for 4 days (Figure 1 - figure supplement 1A). In line with our previous observations, BODIPY staining (Figure 1 - figure supplement 1B), and triglyceride (TG) quantification (Figure 1 - figure supplement 1C) showed a pronounced increase in lipid accumulation after 4 days, without affecting cell viability (Figure 1 - figure supplement 1D).

To assess whether chronic lipid exposure affects BECs *in vivo*, we fed C57BL/6JRj mice for 15 weeks with CD or HFD (Figure 1G) and analyzed their bile ducts. As expected, HFD-fed mice gained weight and developed liver steatosis (Figure 1 - figure supplement 1E), but no apparent fibrosis (data not shown). Of note, HFD-feeding led to an accumulation of lipid droplets in the periportal zone (Figure 1 - figure supplement 1F) and within bile ducts, as reflected by colocalization of BODIPY with PANCK, a BEC marker (Figure 1H), without inducing epithelial damage (Figure 1 - figure supplements 1G-H). Together, these *in vitro* and *in vivo* results demonstrate that BECs accumulate lipids upon chronic FA exposure, raising the question of the functional consequences of this previously unrecognized event on BEC behavior.

### HFD feeding promotes BEC activation and increases organoid formation capacity

To characterize *in vivo* the impact of chronic lipid overload on BECs at the molecular level, we isolated BECs from livers of CD/HFD-fed mice using EPCAM, a pan-BEC marker (Aloia et al., 2019; Pepe-Mooney et al., 2019; Planas-Paz et al., 2019), by fluorescence-activated cell sorting (FACS) (Figure 2A and Figure 2 - supplement 1A). Performing bulk RNA sequencing (RNA-seq) on EPCAM^+^ BECs revealed a diet-dependent clustering in Principal Component Analysis (Figure 2 – supplement 1B), indicating that HFD feeding induces considerable transcriptional changes in BECs *in vivo*. Differential expression analysis further revealed a total of 495 significantly changed genes, 121 upregulated and 374 downregulated (Figure 2 - supplement 1C and Appendix 1 - Table 1). At the same time, HFD promoted the upregulation of *Ncam1* (Figure 2 - supplement 1D), a well-established mediator of BEC activation.

**Figure 2.**
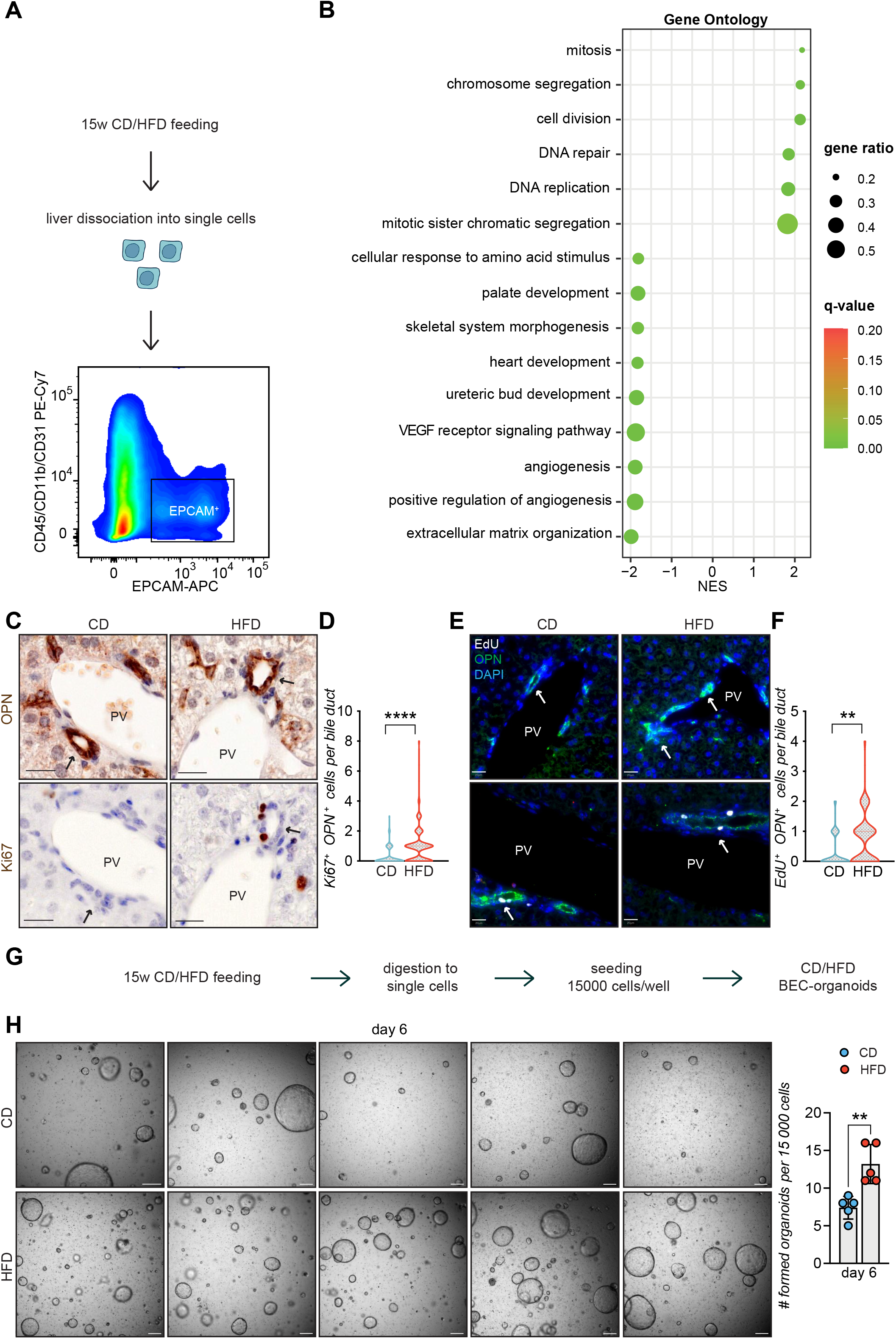
HFD feeding induces EPCAM^+^ BEC proliferation. **(A)** Scheme depicting the isolation of EPCAM^+^ BECs from CD- and HFD-fed mice by FACS. **(B)** Gene set enrichment analysis (GSEA) of Gene Ontology (GO) terms. Top 15 upregulated biological processes (BP), ordered by normalized enrichment score (NES). q-value: false discovery rate adjusted p-values. NES: normalized enrichment score. **(C-F)** Representative co-staining images **(C, E)** and quantification **(D, F)** of BECs stained for OPN and Ki67 **(C-D)**, and OPN and EdU **(E-F)** in livers of CD/HFD-fed mice. n=10 for Ki67 and n=5 for EdU. **(G)** Schematic depicting BEC-organoid formation *in vitro* from CD/HFD-fed mouse livers. **(H)** Images of organoid colonies formed 6 days after seeding, and quantification of organoids per well. n=5. Violin graphs depict the distribution of data points i.e the width of the shaded area represents the proportion of data located there. Other data are shown as mean ± SD. **p < 0.01; ****p < 0.0001; unpaired, two-tailed Student’s t-test was used. PV, portal vein. Arrowheads mark bile ducts. Scale bars, 20 μm (C, E), 200 μm (H). The following figure supplements are available for figure 2: Figure supplement 1. RNA-seq analysis of EPCAM^+^ BECs upon HFD.

To further explore transcriptional changes, we performed gene set enrichment analysis (GSEA) on Gene Ontology (GO) terms (Figure 2B) and KEGG (Figure 2 - supplement 1E) pathways and identified cell proliferation, the most prevalent feature of BEC activation (Sato et al., 2018), as the major upregulated process in these cells upon HFD feeding. Expansion of the reactive BECs requires detachment from their niche and invasion of the parenchyma toward the damaged hepatic area. This process is made possible by reorganizing the extracellular matrix (ECM) and reducing focal adhesion, effectively downregulated in EPCAM^+^ BECs upon HFD (Figure 2B and Figure 2 - supplement 1E).

To validate the RNAseq data, we monitored the activation of BECs *in vivo* by measuring the number of proliferating BECs in the portal region of the livers of mice fed either CD or HFD (Figures 2C-D). Of note, we found that HFD feeding was sufficient to induce a marked increase in the number of active BECs (i.e., Ki67^+^/OPN^+^ cells- Figure 2D). Similar results were observed in an independent cohort of mice challenged with HFD and injected with EdU to track proliferating cells (Figures 2E-F), confirming that chronic lipid exposure stimulates the appearance of reactive BECs within the bile ducts.

The efficiency of BECs to generate organoids *in vitro* has been shown to mirror their regenerative capacity (Aloia et al., 2019). To functionally assess the impact of lipid overload on this process, we measured the organoid forming capacity of isolated BECs, as a read-out of their regenerative functions. To this aim, we quantified the organoid formation efficiency of BECs isolated from CD- and HFD-fed mouse livers (Figure 2G). Strikingly, we observed that HFD-derived BECs were significantly more efficient in generating organoids than their CD counterparts (Figure 2H). Altogether, these results demonstrate that HFD feeding is sufficient to induce, *in vivo*, the exit of BECs from a quiescent state and the acquisition of both proliferative and pro-regenerative features.

### HFD feeding initiates BEC activation via E2Fs

To understand whether the mechanisms underlying BEC activation upon HFD *in vivo* involve canonical processes found in chronically damaged livers, we compared the transcriptional profile of BECs upon HFD with those of DDC-activated BECs (Pepe-Mooney et al., 2019; GSE125688). We identified the most pronounced changes shared between HFD and DDC samples by overlapping separate over-representation enrichment analyses (Appendix 1 - Table 2). Cell division, mitosis, and chromosome segregation were the shared enriched pathways for upregulated genes in HFD and DDC samples (Figure 3A), while ECM organization was the shared enriched pathway for the downregulated genes in HFD and DDC conditions (Figure 3 - supplement 1A). We concluded that the mechanisms of BEC activation induced by lipid overload overlap with those induced by biliary epithelial damage.

**Figure 3.**
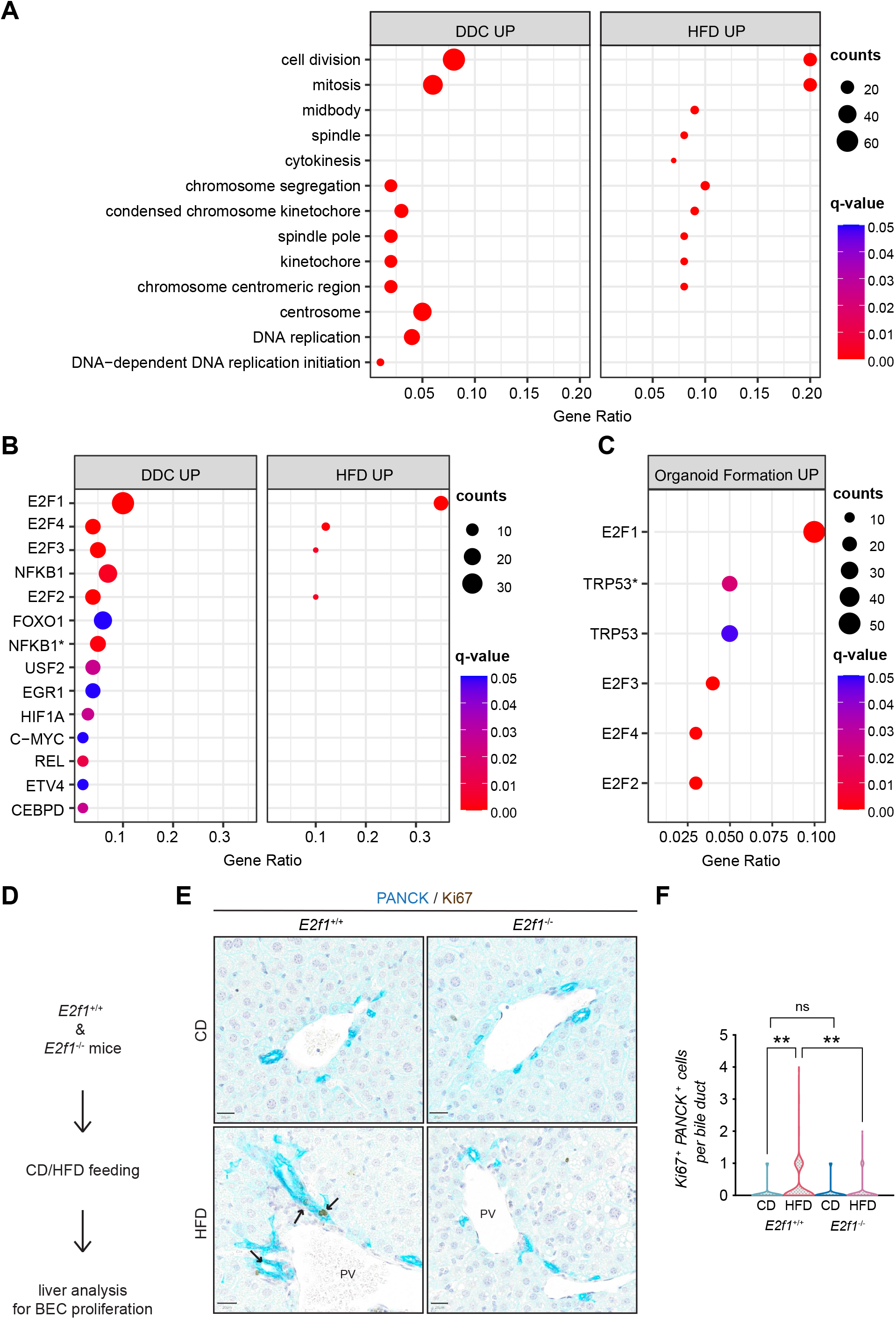
E2Fs are enriched in DDC and HFD datasets and mediate DR initiation *in vivo*. **(A)** Over-representation analysis results. Top 13 enriched biological processes (BP) upon HFD (own data) and DDC (GSE125688) treatment. q-value: false discovery rate adjusted p-values, counts: number of found genes within a given gene set. **(B-C)** Enriched transcription factors (TFs) of upregulated genes identified by over-representation analysis in HFD (own data) and DDC (GSE125688) treatment **(B)**, and during the process of organoid formation from single BECs (Organoids vs T0) (GSE123133) **(C)**. Asterisk (*) marks TFs of the “TF_ZHAO” gene set. **(D)** Schematic depicting *in vivo* E2F1 analysis. **(E-F)** Representative images of PANCK/Ki67 co-staining in livers of *E2f1*^+/+^ and *E2f1*^-/-^ mice fed with CD or HFD **(E)** and quantification of proliferative BECs in the indicated mice **(F)**. For CD, n=5 for *E2f1*^+/+^ and *E2f1*^-/-^. For HFD, n=7 for *E2f1*^+/+^, and n=8 for *E2f1*^-/-^. Violin graphs depict the distribution of data points i.e the width of the shaded area represents the proportion of data located there. ns, not significant; **p < 0.01; two-way ANOVA with Tukey’s test was used. PV, portal vein. Arrowheads mark bile ducts. Scale bars, 20 μm (E). The following figure supplements are available for figure 3: Figure supplement 1. Extended analysis of BECs upon HFD, DDC, and during BEC-organoid formation.

Of note, a more detailed analysis of DDC- and HFD-derived BECs, revealed the concomitant enrichment of 4 overlapping transcription factor (TF) gene sets, E2F1-4 (Figure 3B) and their target genes (Figure 3 - supplement 1B), which have not been linked to DR previously. Moreover, we identified an enrichment of E2Fs (Figure 3C and Appendix 1 - Table 2) and cell division pathway (Figure 3 - supplement 1C) as the most upregulated genes in proliferating BEC-organoids, further corroborating the role of E2Fs in these two *in vitro* (Aloia et al., 2019; GSE123133) and *in vivo* (Pepe-Mooney et al., 2019; GSE125688) DR models.

E2Fs are a large family of TFs with complex functions in cell cycle progression, DNA replication, repair, and G2/M checkpoints (Dimova and Dyson, 2005; Dyson, 1998, 2016; Ren et al., 2002). Therefore, we hypothesized that activation of E2Fs might represent a key event in the process of BEC activation, which is necessary for exiting the quiescent state and driving DR initiation. To test this hypothesis, we focused on E2F1, as it was the most enriched TF in our analysis, and assessed its role in BECs by feeding *E2f1*^+/+^ and *E2f1*^-/-^ mice with HFD (Figure 3D). Remarkably, *E2f1*^-/-^ mice were refractory to BEC activation induced by lipid overload upon HFD, as opposed to *E2f1*^+/+^ mice (Figures 3E-F). These results *in vivo* demonstrate a previously unrecognized role of E2F1 in controlling BEC activation during HFD-induced hepatic steatosis.

### E2F promotes BEC expansion by upregulating glycolysis

The exit of terminally-differentiated cells from their quiescent state requires both energy and building block availability to support cell proliferation. Proliferative cells, therefore, reprogram their glucose metabolism to meet their increased need for biomass and energy (Vander Heiden et al., 2009). Supporting this notion, our interrogation of *in vitro* BEC-organoid formation dataset (Aloia et al., 2019; GSE123133) revealed the enrichment of purine and pyrimidine metabolism, as well as pentose-phosphate pathway, which are tightly connected to glycolysis (Figure 4 - supplement 1A). In line with these findings, a substrate oxidation test in BEC-organoids revealed a preference for glucose, as reflected by the decrease in maximal respiration, when UK5099, a mitochondrial pyruvate carrier inhibitor, was used (Figures 4A-B), while no changes were observed with inhibitors of glutamine (BPTES) and FA (Etomoxir) metabolism (Figure 4 - supplement 1B and 1C).

**Figure 4.**
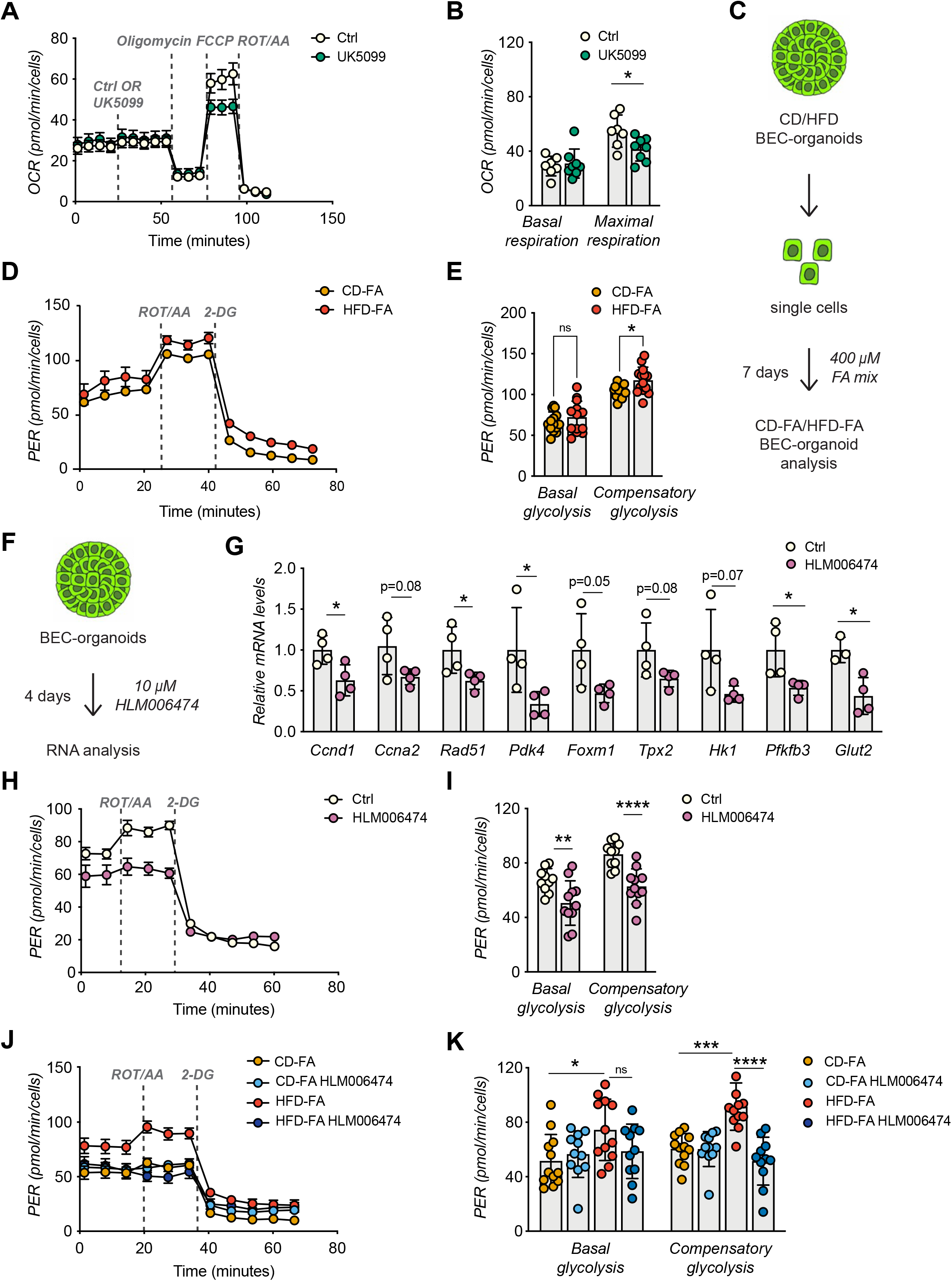
E2Fs promote glycolysis in BEC-organoids. **(A-B)** Seahorse Substrate Oxidation Assay using UK5099, a mitochondrial pyruvate carrier inhibitor **(A)**, and assessment of the glucose dependency **(B)** in CD-derived BEC organoids. n=7 for control (Ctrl), n=8 for UK5099. **(C)** Scheme depicting the treatment of CD/HFD-derived BEC-organoids with FA mix. **(D-E)** Proton efflux rate (PER) **(D)**, and basal and compensatory glycolysis **(E)** measured using Seahorse XF Glycolytic Rate Assay. Relative to panel C. n=13. **(F)** Scheme depicting the treatment of CD-derived BEC-organoids with E2F inhibitor, HLM006474. **(G)** RT-qPCR of selected cell cycle and glycolytic genes, relative to panel H. n=4. (**H-I)** PER during the Seahorse XF Glycolytic Rate Assay **(H)**, and basal and compensatory glycolysis **(I)**, relative to panel F. n=10 for control (Ctrl), n=11 for HLM006474. (**J-K)** PER during the Seahorse XF Glycolytic Rate Assay **(J)**, and basal and compensatory glycolysis **(K)**, relative to panel C and treatment with HLM006474. n=12 for control (Ctrl), n=11 for HLM006474. Data are shown as mean ± SD (SEM for A, D, H, J). Absence of stars or ns, not significant (p > 0.05); *p < 0.05; **p < 0.01; ***p <0.001; ****p < 0.0001; unpaired, two-tailed Student’s t-test (G), two-way ANOVA with Sidak’s test (B, E, I, K) were used. The following figure supplements are available for figure 4: Figure supplement 1. E2F activation correlates with increased glycolysis.

To investigate the metabolic changes in BEC-organoids upon HFD, we treated CD/HFD BEC-organoids with FA mix to mimic steatotic conditions *in vitro* (Figure 4C). We hypothesized that the presence of glucose and FA in culture media would reveal a metabolic shift of BEC-organoids. Consistent with our hypothesis, HFD-FA BEC-organoids demonstrated increased compensatory glycolytic rates (Figures 4D-E and Figure 4 - supplement 1D). Of note, there was a reduction in oxidative phosphorylation in HFD-FA BEC-organoids, as evidenced by the decrease in maximal respiration (Figure 4 - supplement 1E-G), which might reflect their preference for glycolytic pathway to generate biomass.

Besides their prominent role in cell cycle progression, E2Fs coordinate several aspects of cellular metabolism (Denechaud et al., 2017; Nicolay and Dyson, 2013), and promote glycolysis in different contexts (Blanchet et al., 2011; Denechaud et al., 2015; Huber et al., 2021). These findings prompted us to postulate that E2F might control glycolysis and thus the glucose preference observed in BEC-organoids. To investigate this hypothesis, we treated BEC-organoids with an E2F inhibitor, HLM006474 (Figure 4F). As expected, HLM006474 treatment reduced the transcriptional levels of several genes involved in cell cycle progression and glycolytic metabolism (Figure 4G), and decreased the glycolytic flux, as evidenced by the blunted proton efflux rate (PER) (Figures 4H-I). Moreover, E2F inhibition was able to reverse the metabolic phenotype only in HFD-FA BEC-organoids (Figures 4J-K).

In conclusion, these results demonstrate that HFD-induced E2F activation controls the conversion of BECs from quiescent to active progenitors by promoting the expression of cell cycle genes while simultaneously driving a shift towards glycolysis.

## Discussion

Through DR activation, BECs represent an essential reservoir of progenitors that are crucial for coordinating hepatic epithelial regeneration in the context of chronic liver diseases (Choi et al., 2014; Deng et al., 2018; Español–Suñer et al., 2012; Huch et al., 2013; Lu et al., 2015; Raven et al., 2017; Rodrigo-Torres et al., 2014; Russell et al., 2019). BEC functions are tightly controlled by YAP metabolic pathways (Meyer et al., 2020; Pepe-Mooney et al., 2019; Planas-Paz et al., 2019) and recent studies from different tissues have provided evidence that specific metabolic states play instructive roles in controlling cell fate and tissue regeneration (Beyaz et al., 2016; Capolupo et al., 2022; Miao et al., 2020; Zhang et al., 2016). Aberrant lipid accumulation is a hallmark of early NAFLD, and imbalances in lipid metabolism are known to affect hepatocyte homeostasis, including induction of lipo-toxicity and cell death (X. Wang et al., 2016; Sano et al., 2021; De Gottardi et al., 2007; Wobser et al., 2009; Ipsen et al., 2018). However, the role of lipid dysregulation in BECs and whether it has an impact on the initiation of DR remains unexplored in the setting of NAFLD.

Here, using HFD-induced mouse models, we studied BEC metabolism in steatosis, the first stage of NAFLD, and demonstrated lipid accumulation in BECs during chronic HFD *in vivo* and their resistance to lipid-induced toxicity. By using BEC-organoids, we observed that FAs directly target BECs, without any involvement of hepatocytes and that BECs functionally respond to lipid overload. Importantly, BEC-organoids derived from CD- and HFD-fed mouse livers were shown to shift their cellular metabolism toward more glycolysis in the presence of lipids. Furthermore, we found that the HFD-induced metabolic shift was sufficient to reprogram BEC identity *in vivo*, allowing their exit from a quiescent state and the simultaneous acquisition of progenitor functions, such as proliferation and organoid-initiating capacity, two hallmarks of DR. These results highlight the metabolic plasticity of BECs and shed light on an unpredicted mechanism of BEC activation in HFD-induced hepatic steatosis. Importantly, we observed that lipid overload is sufficient to induce BEC activation in steatotic livers and that this process precedes parenchymal damage, thus resolving a long-lasting debate on the chronology of DR initiation in chronic liver diseases. While the functional contribution of lipid-activated BECs in liver regeneration during the late stages of NAFLD will require further studies, our data clearly point out the role of BECs as sensors, and possibly effectors, of early liver diseases such as steatosis.

To fully understand the underlying basis of the observed phenotype, we characterized the transcriptome of primary BECs derived from steatotic livers of HFD-fed mice and demonstrated that long-term feeding of a lipid-enriched diet strongly promotes BEC proliferation, suggesting a strong link between metabolic adaptation and progenitor function. By combining our transcriptomic analysis with data mining of publicly available DR datasets (Aloia et al., 2019; Pepe-Mooney et al., 2019), we identified the E2F transcription factors as master regulators of DR in the context of NAFLD. Moreover, the expression of *Pdk4,* an E2F1 target (Hsieh et al., 2008; L.-Y. Wang et al., 2016), was upregulated in FA-treated BEC-organoids and upon HFD. PDK4 limits the utilization of pyruvate for oxidative metabolism while enhancing glycolysis, which reinforces our data demonstrating that E2Fs rewire BEC metabolism toward glycolysis to fuel progenitor proliferation. These observations feature E2Fs as the molecular rheostat integrating the metabolic cell state with the cell cycle machinery to coordinate BEC activation. However, how E2Fs are regulated upon HFD and whether they are interconnected with the already known YAP, mTORC1, and TET1 pathways remain unknown and will require further investigations.

While our data are derived from obese mice, a recent report showed increased numbers of cholangiocytes in steatotic human livers (Hallett et al., 2022) and E2F1 has been found to be upregulated in the livers of obese patients (Denechaud et al., 2015). Moreover, human subjects with elevated visceral fat demonstrated increased glucose metabolism (Broadfield et al., 2021). These observations, while correlative, set the ground for future research in understanding the role and the therapeutic potential of lipid metabolism and E2Fs in controlling BEC activation, thus initiating DR in humans.

## Materials and Methods

### Mouse studies and ethical approval

All the animal experiments were authorized by the Veterinary Office of the Canton of Vaud, Switzerland under license authorization VD3721 and VD2627.b. C57BL/6JRj mice were obtained from Janvier Labs and *E2f1^+/+^* and *E2f1^−/−^* (B6;129S4-E2f1tm1 Meg/J) mice were purchased from The Jackson Laboratory. 8-week-old C57BL/6JRj male mice were fed with Chow Diet (CD - SAFE Diets, SAFE 150) or High Fat Diet (HFD - Research Diets Inc, D12492i) for 15 weeks. 7-week-old *E2f1^+/+^* and *E2f1^−/−^* male mice were fed with Chow Diet (CD – Kliba Nafag 3336) or High Fat Diet (HFD - Envigo, TD93075) for 29 weeks. The well-being of the animals was monitored daily, and body weight was monitored once per week until the end of the experiment. All mice had unrestricted access to water and food and liver tissues were harvested at the end of the experiment.

### Data reporting

Mice were randomized into different groups according to their genotype. A previous HFD experiment was used to calculate the sample size for C57BL/6JRj mouse experiments. Mice showing any sign of severity, predefined by the Veterinary Office of the Canton of Vaud, Switzerland were sacrificed and excluded from the data analyses. *In vitro* experiments were repeated with at least 3 biological replicates (BEC-organoids from different mice) or were repeated at least twice by pooling 4 mice per condition (for Seahorse analysis).

### Proliferation Assay

Cell proliferation was assessed by EdU assay (Click-iT EdU Alexa Fluor 647, ThermoFisher, C10340) following the manufacturer’s instructions. For *in vivo* studies, EdU was resuspended in phosphate-buffered saline (PBS- ThermoFisher, 10010002) and 200 μL of the solution was injected intraperitoneally (50 μg per g of mouse weight) 16 hours before the sacrifice.

### EPCAM^+^ BEC isolation and FACS analysis

23-week-old CD/HFD-fed C57BL/6JRj male mice were used for this experiment and sacrificed in the fed state. To isolate the BECs, mouse livers were harvested and digested enzymatically as previously reported (Broutier et al., 2016). Briefly, livers were minced and incubated in a digestion solution (1% fetal bovine serum (FBS) (Merck/Sigma, F7524) in DMEM/Glutamax (ThermoFisher, 31966-021) supplemented with HEPES (ThermoFisher, 15630-056) and Penicillin/Streptomycin (ThermoFisher, 15140-122) containing 0.0125% (mg/ml) collagenase (Merck/Sigma, C9407), 0.0125% (mg/ml) dispase II (ThermoFisher, 17105-041) and 0.1 mg/ml of DNAase (Merck/Sigma, DN25). This incubation lasted 2-3h on a shaker at 37°C at 150 rpm. Livers were then dissociated into single cells with TrypLE (GIBCO, 12605028) and washed with washing buffer (1% FBS (Merck/Sigma, F7524) in Advanced DMEM/F-12 (GIBCO, 12634010) supplemented with Glutamax (ThermoFisher, 35050061), HEPES (ThermoFisher, 15630-056) and Penicillin/Streptomycin (ThermoFisher, 15140-122)). Single cells were filtered with a 40 µm cell strainer (Falcon, 352340) and incubated with fluorophore-conjugated antibodies CD45–PE/Cy7 (BD Biosciences, 552848), CD11b– PE/Cy7 (BD Biosciences, 552850), CD31–PE/Cy7 (Abcam, ab46733) and EPCAM–APC (eBioscience, 17-5791-82) for 30 min on ice. BECs were sorted using FACSAria Fusion (BD Biosciences) as previously described (Aloia et al., 2019). Briefly, individual cells were sequentially gated based on cell size (forward scatter (FSC) versus side scatter (SSC)) and singlets. BECs were then selected based on EPCAM positivity after excluding leukocytes (CD45^+^), myeloid cells (CD11b^+^), and endothelial cells (CD31^+^), yielding a population of single CD45^-^/CD11b^-^/ CD31^-^/EPCAM^+^ cells. All flow cytometry data were analyzed with FlowJo v10.8 software (BD Life Sciences).

### RNA preparation from EPCAM^+^ BECs and bulk RNA-seq data analysis

RNA was isolated from sorted BECs using the RNeasy micro kit (QIAGEN, 74104) and the amount and quality of RNA were measured with the Agilent Tapestation 4200 (Agilent Technologies, 5067-1511). As a result, RNA-seq of 5 CD and 7 HFD samples was performed by BGI with the BGISEQ-500 platform. FastQC was used to verify the quality of the reads (Andrews, 2010). No low-quality reads were present, and no trimming was needed. Alignment was performed against the mouse genome (GRCm38) following the STAR (version 2.6.0a) manual guidelines (Dobin et al., 2013). The obtained STAR gene counts for each alignment were analyzed for differentially expressed genes using the R package DESeq2 (version 1.34.0) (Love et al., 2014). A threshold of 1 log2 fold change and adjusted p-value smaller than 0.05 were considered when identifying the differentially expressed genes. A principal component analysis (PCA) (Lê et al., 2008) was used to explore the variability between the different samples.

### Gene set enrichment analysis (GSEA)

We used the clusterProfiler R package (Yu et al., 2012) to conduct GSEA analysis on various gene sets. Gene sets were retrieved from http://ge-lab.org/gskb/ for *M.musculus*. We ordered the differentially expressed gene list by log2 (Fold-changes) for the analysis with default parameters.

### Over-representation enrichment analysis

All significantly changing genes (adjusted p-value < 0.05 and an absolute fold change > 1) were split into 2 groups based on the direction of the fold change (genes significantly up- & down-regulated). An over-representation analysis using the clusterProfiler R package was performed on each of the two groups to identify biologically overrepresented terms.

### Figure generation with R

The R packages ggplot2 (Wickham, 2016) retrieved from https://ggplot2.tidyverse.org and ggpubr were used to generate figures.

### Culture of mouse liver BEC-organoids from biliary duct fragments and single cells

BEC-organoids were established from bile ducts of C57BL/6JRj male mice as previously described (Broutier et al., 2016; Sorrentino et al., 2020). Thus, the liver was digested as detailed above (EPCAM^+^ BEC isolation) and bile ducts were isolated, they were pelleted by centrifugation at 200 rpm for 5 min at 4°C and washed with PBS twice. Isolated ducts were resuspended in Matrigel (Corning, 356231) and cast in 10 µl droplets in 48-well plates. When gels were formed, 250 µl of isolation medium (IM-Advanced DMEM/F-12-Gibco,12634010) supplemented with Glutamax (ThermoFisher, 35050061), HEPES (ThermoFisher, 15630-056), and Penicillin/Streptomycin (ThermoFisher, 15140-122), 1X B27 (Gibco, 17504044), 1mM *N*-acetylcysteine (Sigma-Aldrich, A9165), 10 nM gastrin (Sigma-Aldrich, G9145), 50 ng/ml EGF (Peprotech, AF-100-15), 1 µg/ml Rspo1 (produced in-house), 100 ng/ml FGF10 (Peprotech, 100-26), 10 mM nicotinamide (Sigma-Aldrich, N0636), 50 ng/ml HGF (Peprotech, 100-39), Noggin (100 ng/ml produced in-house), 1µg/ml Wnt3a (Peprotech, 315-20) and 10 μM Y-27632 (Sigma, Y0503) was added to each well. Plasmids for Rspo1 and Noggin production were a kind gift from Joerg Huelsken. After the first 4 days, IM was replaced with the expansion medium (EM), which was the IM without Noggin, Wnt3a, and Y-27632. For passaging, organoids were removed from Matrigel a maximum of one week after seeding and dissociated into single cells using TrypLE Express (Gibco, 12604013). Single cells were then transferred to fresh Matrigel. Passaging was performed in a 1:3 split ratio.

For the FA-treatment of BEC-organoids, palmitic acid (Sigma, P0500) and oleic acid (Sigma, O1008) were dissolved in 100% ethanol into 500 and 800 μM stock solutions respectively, and kept at −20 °C. For each experiment, palmitic acid and oleic acid were conjugated to 1% fatty acid free bovine serum albumin (BSA) (Sigma, A7030), in EM through 1:2000 dilution each (Malhi et al., 2006). The concentration of vehicle, ethanol, was 0.1% ethanol in final incubations and 1% fatty acid free BSA in EM was used as the control for FA treatment.

### Liver immunohistochemistry (IHC) and immunofluorescence (IF)

For liver paraffin histology, livers were washed in PBS (Gibco, 10010023), diced with a razor blade, and fixed overnight in 10% formalin (ThermoFisher, 9990244) while shaking at 4°C. The next day fixed livers were washed twice with PBS, dehydrated in ascending ethanol steps, followed by xylene, and embedded in paraffin blocks. 4 μm thick sections were cut from paraffin blocks, dewaxed, rehydrated, and quenched with 3% H_2_O_2_ for 10 minutes to block the endogenous peroxidase activity (for IHC). Antigen retrieval was performed by incubating the sections in 10 mM citrate buffer (pH 6.0) for 20 min at 95 °C. After the sections cooled to room temperature, they were washed and blocked with blocking buffer (1% BSA (Sigma, A7906) and 0.5% Triton X-100 (Sigma, X100) in PBS), for 1 h at room temperature. The primary antibodies anti-Ki67 (ThermoFisher, MA5-14520), anti-PANCK (Novusbio, NBP600-579), anti-OPN (R&D Systems, AF808), anti-Cleaved caspase-3 (Cell Signaling, 9661) were diluted in a 1:100 dilution of the blocking buffer and incubated overnight at 4 °C. For IHC, ImmpRESS HRP conjugated secondary (VectorLabs MP-74-01-15 and MP-74-02) were incubated for 30 min and detection was performed by using a 3.3’-diaminobenzidine (DAB) reaction. Sections were counterstained with Harris and mounted. For IF, sections were washed and incubated for 1 h with Alexa Fluor conjugated secondary antibodies (1:1000 in blocking solution; Invitrogen). Following extensive washing, sections were counterstained with DAPI (ThermoFisher, 62248), and mounted in ProLong Gold Antifade Mountant (Thermo Fischer, P36930).

For IF of liver cryosections, the livers were frozen in O.C.T. compound (VWR chemicals) on dry ice-filled with isopentane. 10 μm liver sections were cut from O.C.T embedded samples, hydrated, and washed twice in PBS. The sections were blocked in blocking buffer for 1h at room temperature and incubated with BODIPY 558/568 (Invitrogen, D38D35) for 20 minutes. After fixation with 4% paraformaldehyde (PFA) solution (Sigma, 1004960700) for 15 minutes, sections were washed with PBS. Then, sections were permeabilized using 5% BSA in TBS-T and stained with primary antibody anti-PANCK diluted in blocking buffer for 16 h at 4 °C. The next day, the sections were washed three times with PBS and the appropriate Alexa Fluor secondary antibodies were diluted in blocking buffer (1:1000) and incubated with the sections for 1 h at room temperature. The sections were washed in PBS and incubated with DAPI diluted 1:1000 in PBS for 1 h at room temperature. Finally, the sections were mounted in ProLong Gold Antifade Mountant.

Stained sections were imaged by a virtual slide microscope (VS120, Olympus) and analysis was performed using QuPath software (Bankhead et al., 2017).

### BEC-organoid whole-mount immunofluorescence

BEC-organoids were incubated with BODIPY 558/568 for 20min, and then washed with PBS and extracted from Matrigel using Cell Recovery Solution (Corning, 354253). After fixing with 4% PFA in PBS (30 min, on ice), they were pelleted by gravity to remove the PFA and were washed with PBS and ultra-pure water. BEC-organoids were then spread on glass slides and allowed to attach by drying. The attached BEC-organoids were rehydrated with PBS and permeabilized with 0.5% Triton X-100 in PBS (1 h, room temperature) and blocked for 1 h in blocking buffer. After washing with PBS, samples were incubated for 1 h at room temperature with Alexa Fluor Phalloidin 488 (Invitrogen, A12379). Following extensive washing, samples were counterstained with DAPI and were imaged by a confocal microscope (LSM 710, Zeiss). Signal intensity was adjusted on each channel using Fiji software (Schindelin et al., 2012).

### Quantitative real-time qPCR for mRNA quantification

BEC-organoids were extracted from Matrigel using Cell Recovery Solution (Corning, 354253). RNA was extracted from organoid pellets using the RNAqueous total RNA isolation kit (Invitrogen, AM1931) and the RNeasy Micro Kit (Qiagen, 74004) following the manufacturer’s instructions. RNA was transcribed to complementary DNA using QuantiTect Reverse Transcription Kit (Qiagen, 205314) following the manufacturer’s instructions. PCR reactions were run on the LightCycler 480 System (Roche) using SYBR Green (Roche, 4887352001) chemistry. Real-time quantitative polymerase chain reaction (RT-qPCR) results were presented relative to the mean of *36b4* (comparative ΔCt method). Primers for RT-qPCR are listed in Appendix 1 - Table 3.

### E2F inhibition

For the E2F inhibition experiment, single BECs were grown for 7 days and allowed to form organoids. For the Seahorse experiment, BEC-organoids were treated with E2F inhibitor, HLM006474 (10 μM, Merck, 324461), overnight before the metabolic assay. For RT-qPCR analysis, BEC-organoids were treated with HLM006474 chronically for 4 days.

### Bioenergetics with Seahorse extracellular flux analyzer

The oxygen consumption rate (OCR), extracellular acidification rate (ECAR), and proton-efflux rate (PER) of the BEC-organoids were analyzed by an XFe96 extracellular flux analyzer (Agilent) following the manufacturer’s instructions according to assay type.

For Mito Stress Test on CD/HFD-derived BEC-organoids, the organoids were grown with FA mix for 7 days. On day 7, 10 μM HLM006474 or DMSO as vehicle were added overnight. The next morning, BEC-organoids were dissociated, and 20000 cells were seeded with Seahorse Assay Medium in XFe96 Cell Culture Microplates (Agilent, 101085-004), which were previously coated with 10% Matrigel in Advanced DMEM/F-12. Seahorse Assay Medium was unbuffered, serum-free pH 7.4 DMEM supplemented with 10 mM glucose (Agilent, 103577-100), 10 mM pyruvate (Gibco, 11360070), and 2 mM glutamine (Agilent, 103579-100), and 10 μM HLM006474 or DMSO (vehicle) were added when indicated. After 2 h incubation for cell attachment, plates were transferred to a non-CO^2^ incubator at 37 °C for 45 minutes. Mitochondrial OCR was measured in a time course before and after the injection of 1.5 μM Oligomycin (Millipore, 495455), 2.5 μM FCCP (Sigma, C2920), and 1 μM Rotenone (Sigma, R8875)/Antimycin A (Sigma, A8674).

For Glycolytic Rate Assay, CD BEC-organoids were grown without FA mix and CD/HFD-derived BEC-organoids were grown with FA mix for 7 days. The Seahorse assay preparations including the E2F inhibitor were the same as mentioned above. GlycoPER was measured in a time course before and after the injection of 1 μM Rotenone/Antimycin A, and 500 μM 2-DG (Sigma, D8375).

For Substrate Oxidation Assay, CD BEC-organoids were grown without FA mix for 7 days. On day 8, they were dissociated and prepared for Seahorse assay, without E2F inhibitor. Mitochondrial OCR was measured in a time course before and after the injection of Oligomycin (1.5 μM), FCCP (2.5 μM), and Rotenone/Antimycin A (1 μM) with or without UK5099 (Sigma, PZ0160), Etomoxir (E1905) and BPTES (SML0601), inhibitors of glucose oxidation, fatty acid oxidation and glutamine oxidation, respectively, in separate experiments.

All Seahorse experiments were normalized by cell number through injection of 10 μM of Hoechst (ThermoFisher,62249) in the last Seahorse injection. Hoechst signal (361/486 nm) was quantified by SpectraMax iD3 microplate reader (Molecular Devices).

### BEC-organoid growth assay

BEC-organoid formation efficiency was quantified by counting the total number of cystic/single layer (lumen-containing) CD/HFD-derived BEC-organoids 6 days after seeding and normalizing it to the total number of cells seeded initially (15000 cells). Organoids were imaged by DM IL LED inverted microscope (Leica), selected as regions of interest (ROI) using widefield 4x magnification, and counted manually.

### BEC-organoid functional analysis

Grown BEC-organoids were treated with the FA-mix for 4 days and triglyceride levels were measured with a Triglyceride kit (Abcam, ab65336) following the manufacturer’s instructions. Cell-titer Glo (Promega, G7570) was used to investigate cell viability. For functional assays involving single BECs, grown organoids were dissociated into single cells. 10000 BECs were seeded, and organoid formation was allowed for 7 days. Cell viability, apoptosis, and cell death were investigated using Cell-titer Glo, Caspase 3/7 activity (Promega, G8091), and Nucgreen Dead 488 staining (Invitrogen, R37109), respectively, according to the protocol of manufacturers. For cell death staining, organoids were imaged using ECLIPSE Ts2 inverted microscope (Nikon).

### Quantification and statistical analysis

Data were presented as mean ± standard deviation (mean ± SD.) unless it is stated otherwise in the figure legend. *n* refers to biological replicates and is represented by the number of dots in the plot or stated in the figure legends. For the Seahorse experiments, *n* refers to technical replicates pooled from 4 biological replicates and is represented by the number of dots in the plot or stated in the figure legends. The statistical analysis of the data from bench experiments was performed using Prism (Prism 9, GraphPad). The differences with p<0.05 were considered statistically significant. No samples (except outliers) or animals were excluded from the analysis. Data are expected to have a normal distribution.

For two groups comparison, data significance was analyzed using a two-tailed, unpaired Student’s t-test. In case of comparisons between more than two groups, one- or two-way ANOVA was used. Dunnet’s, Tukey’s, or Sidak’s tests were used to correct for multiple comparisons. Statistical details of each experiment can be found in the respective figure legends.

### Data availability

Computational analysis was performed using established packages mentioned in previous sections, and no new code was generated. Two publicly available RNA-Seq datasets of mouse BECs with accession numbers GSE123133 (Aloia et al., 2019) and GSE125688 (Pepe-Mooney et al., 2019) were downloaded from the GEO and used for GSEA and over-representation enrichment analysis as mentioned previously.

## Acknowledgments

We thank the platforms of the SV faculty for support, Sabrina Bichet, Fabiana Fraga, and Jéromine Imbach for technical assistance, and Yu Sun for the critical review of manuscript figures. This work was funded by the Kristian Gerhard Jebsen Foundation and the Ecole Polytechnique Fédérale de Lausanne (EPFL).

## Author contributions

E.Y. and K.S. conceived and designed the project. G.S. and A.P. helped with experiment planning and manuscript preparation. E.Y. performed the experiments, analyzed the data, and wrote the original manuscript draft. G.E.A. performed bioinformatics analysis under the supervision of J.A.. A.J. generated the FACS figure. P.D.D. and K.H. provided the *E2f1*^+/+^ and *E2f1*^-/-^ liver samples under the supervision of L.F.. K.S. supervised the work. The manuscript was edited by all co-authors.

## Declaration of interests

The authors declare no competing interests.

## Figure Supplements and Legends

**Figure 1 - figure supplement 1:**
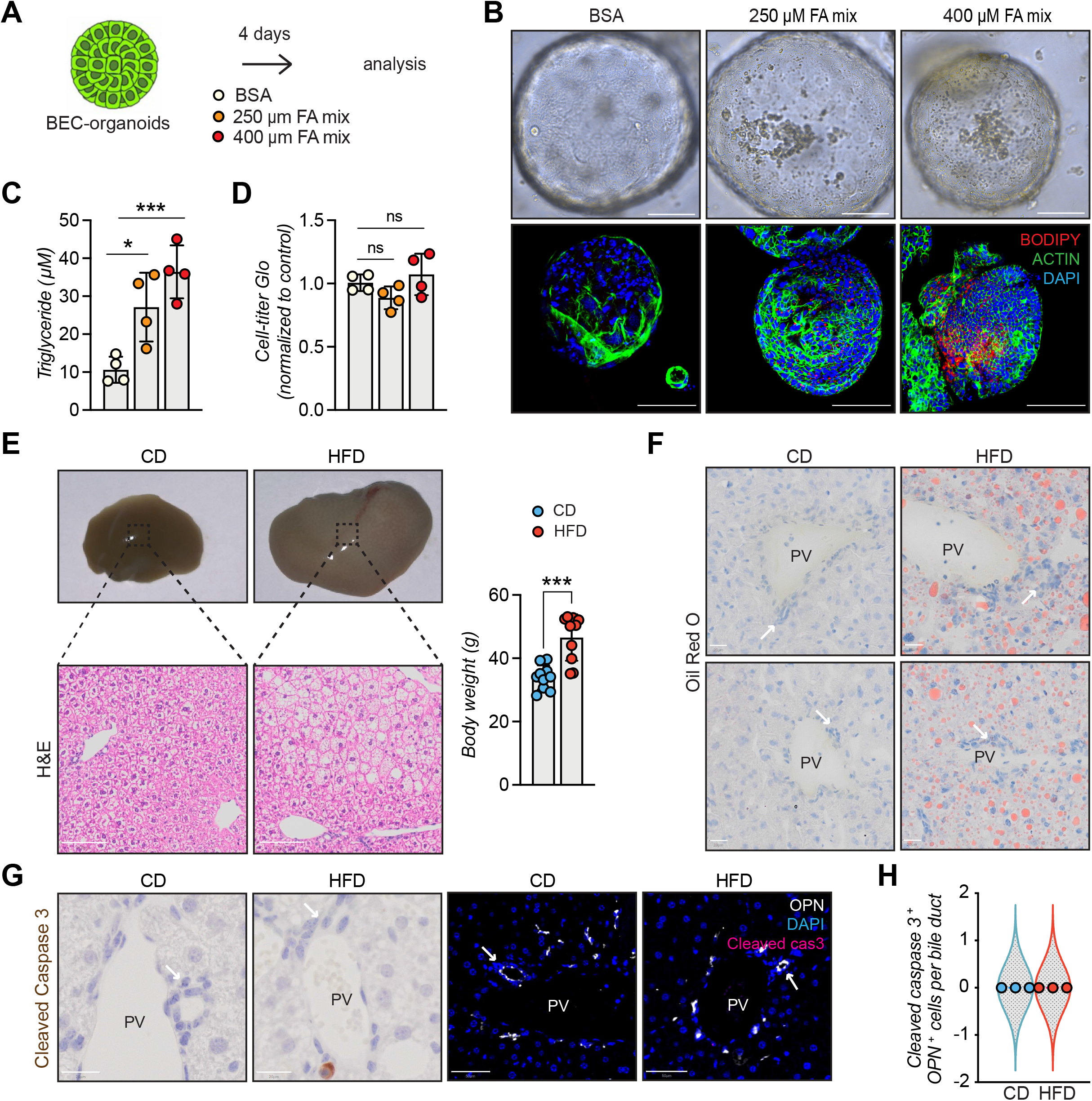
Further characterization of lipid accumulation in BECs. **(A)** Scheme depicting FA mix treatment for already grown BEC-organoids for 4 days. **(B)** Representative brightfield and immunofluorescence (IF) images of lipids (BODIPY) and ACTIN co-staining, relative to panel A. n=4. **(C)** Measurement of triglyceride concentration upon FA mix treatment relative to panel A, normalized by protein amount. n=4. **(D)** Cell-titer Glo measurement for viability detection relative to panel A. n=4. **(E)** Representative images of CD/HFD-fed mouse livers and hematoxylin and eosin (H&E) staining and the final body-weight measurement at the end of the experiment. n=10. **(F)** Representative Oil Red O staining, relative to panel E. n=8. **(G- H)** Representative cleaved caspase 3 stainings, relative to panel E **(G)** and quantification of apoptotic cells in bile ducts **(H)**. n=3. Data are shown as mean ± SD. Absence of stars or ns, not significant (p > 0.05); *p < 0.05; ***p < 0.001; one-way ANOVA with Dunnet’s test (C, D) or unpaired, two-tailed Student’s t-test (E) were used. PV, portal vein. Arrowheads mark bile ducts. Scale bars, 100 μm (B-E), 50 μm (G-IF), and 20 μm (F, G-brightfield).

**Figure 2 - figure supplement 1:**
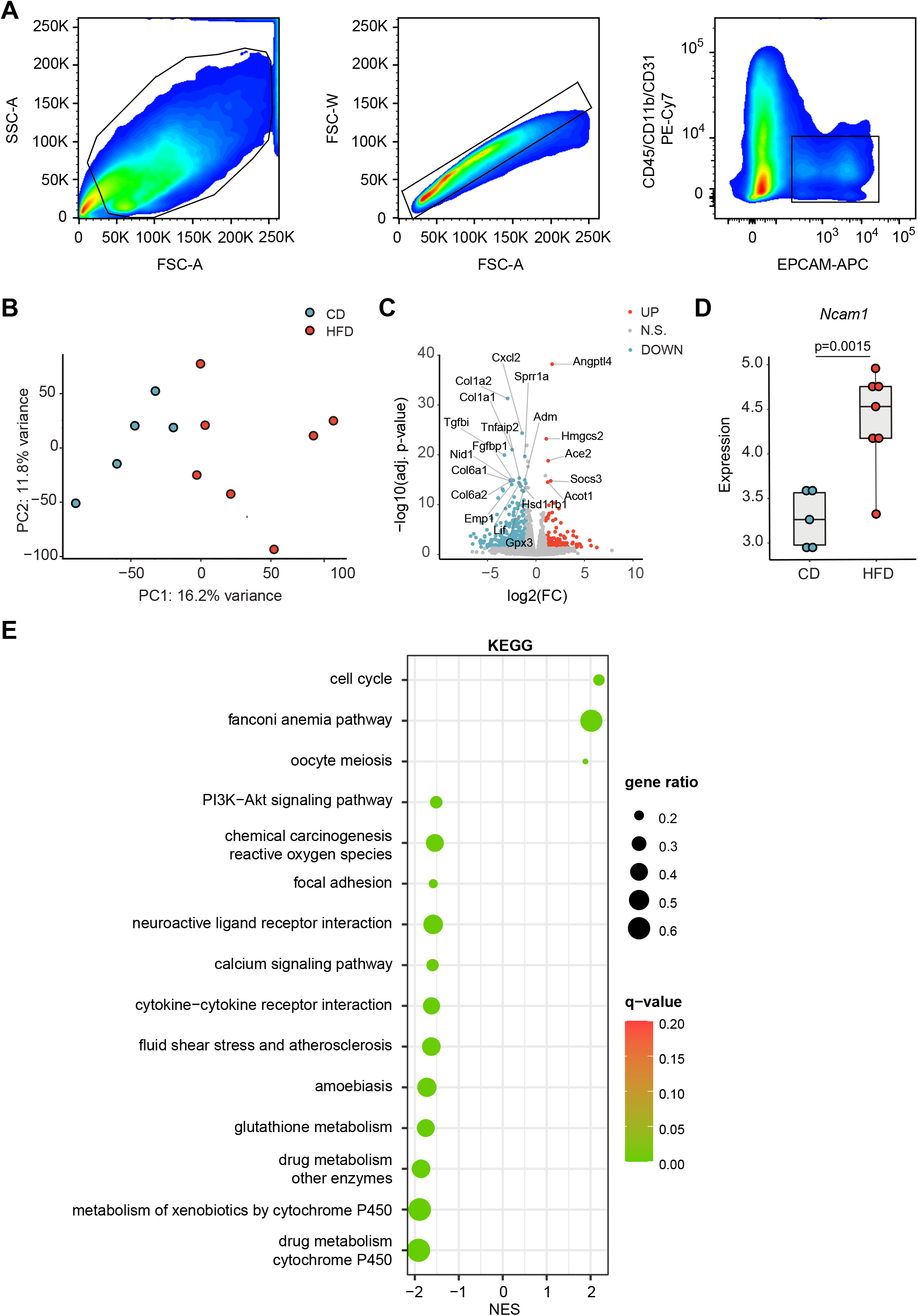
RNA-seq analysis of EPCAM^+^ BECs upon HFD. **(A)** FACS gating strategy for isolation of CD45^−^/CD11b^−^/CD31^−^/EpCAM^+^ BECs: individual cells were sequentially gated based on cell size (FSC versus SSC) and singlets. BECs were then selected based on EPCAM positivity after excluding leukocytes (CD45^+^), myeloid cells (CD11b^+^), and endothelial cells (CD31^+^), yielding a population of single CD45^-^/CD11b^-^/ CD31^-^/EPCAM^+^ BECs. **(B)** Principal component analysis (PCA) of mRNAs measured in mice fed CD or HFD by RNA-seq. n= 5 for CD, n=7 for HFD. **(C)** Volcano plot of HFD vs CD differential analysis. Top 20 differentially expressed genes were labeled. Blue dots represent downregulated genes (log2(FC) < −1 & adj. p-value < 0.05). Red dots represent upregulated genes (log2(FC) > 1 & adj. p-value < 0.05). Grey dots represent genes not changing significantly. **(D)** Box plot representing the differential gene expression of *Ncam1*. n= 5 for CD, n=7 for HFD. The Y-axis depicts log2(cpm+1) values. **(E)** Gene set enrichment analysis (GSEA) of KEGG terms. Top 15 enriched pathways (sorted by q-value). q-value: false discovery rate adjusted p-values. NES: normalized enrichment score. Data are shown as mean ± SD. Unpaired two-tailed Student’s t-test was used.

**Figure 3 - figure supplement 1:**
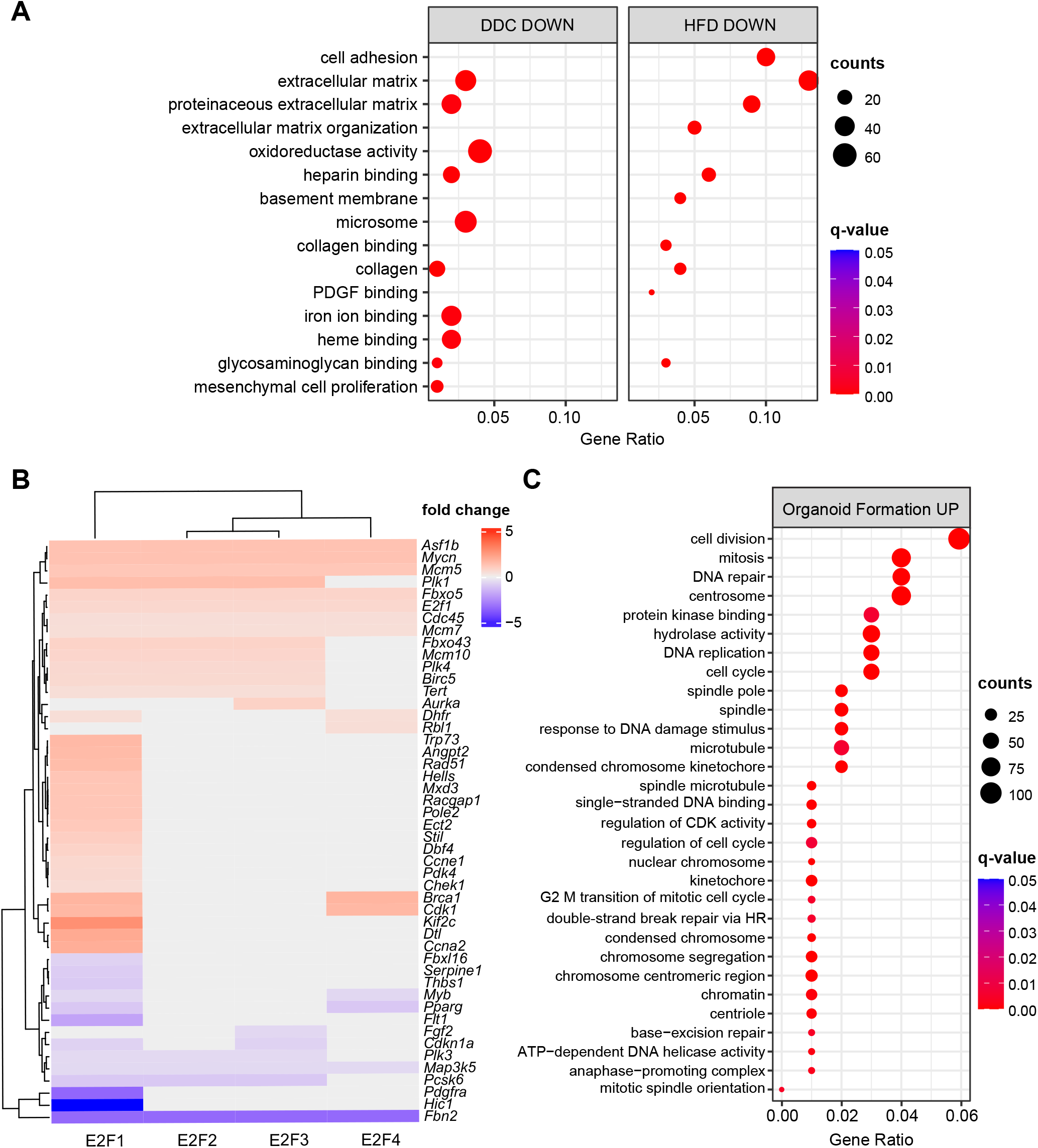
Extended analysis of BECs upon HFD, DDC, and during BEC-organoid formation. **(A)** Over-representation analysis results. Top 15 enriched biological processes (BP) upon HFD (own data) and DDC (GSE125688) treatment in BECs. q-value: false discovery rate adjusted p-values, counts: number of found genes within a given gene set. **(B)** Heatmap of E2F1-4 target genes from TF gene sets. Genes with absolute fold change lower than 0.5 were not shown. **(C)** GO over-representation analysis of upregulated BP during the process of organoid formation from single BECs (Organoids vs T0) (GSE123133).

**Figure 4 - figure supplement 1:**
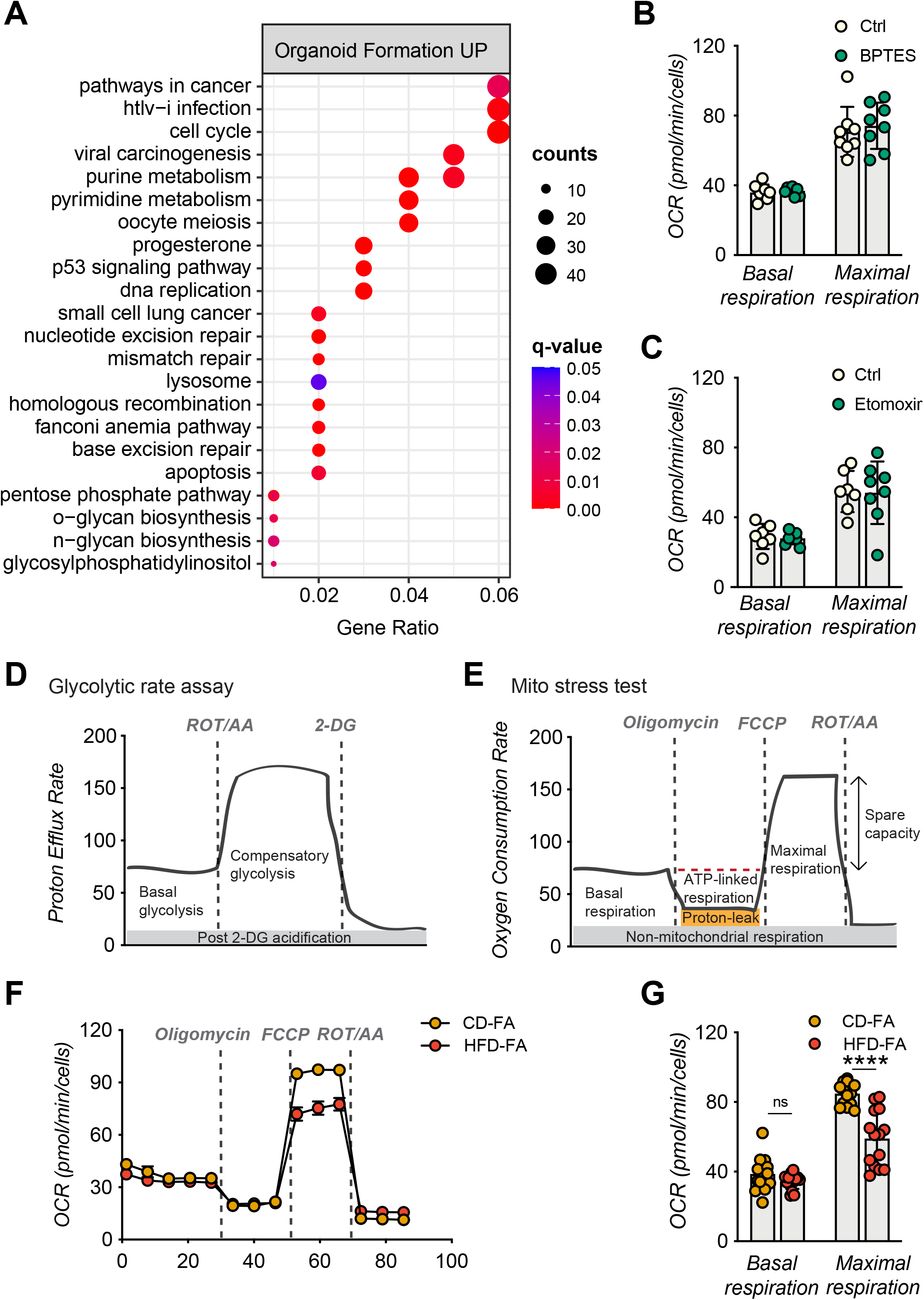
E2F activation correlates with increased glycolysis. **(A)** Over-representation analysis results. Top 22 enriched KEGG and EHMN pathways during the process of organoid formation from single BECs (Organoids vs T0) (GSE123133). q-value: false discovery rate adjusted p-values, counts: number of found genes within a given gene set. **(B-C)** Seahorse Substrate Oxidation Assay using BPTES, n=8 **(B)**, or Etomoxir, n=7 for control (Ctrl) and n=8 for inhibitor conditions **(C)** in CD-derived BEC-organoids. **(D-E)** Scheme depicting Seahorse Glycolytic Rate Assay **(D),** and Mito Stress Test **(E)**. **(F-G)** Oxygen consumption rate (OCR) **(F)**, and basal and maximal respiration **(G)** measured using Seahorse XF Mito Stress Test. Relative to panel C. n=14. Data are shown as mean ± SD (SEM for F). absence of stars or ns, non-significant (p-value > 0.05); ****p < 0.0001; two-way ANOVA with Sidak’s test (B, C, G) was used.

## Table Supplements and Legends

**Appendix 1 - Table 1. Differential expression analysis results of EPCAM^+^ BECs upon HFD. Related to Figure 2.**

**Appendix 1 - Table 2. Over-representation analysis of upregulated and downregulated genes in HFD, DDC, and BEC-organoid formation datasets. Related to Figure 3.**

**Appendix 1 - Table 3.**
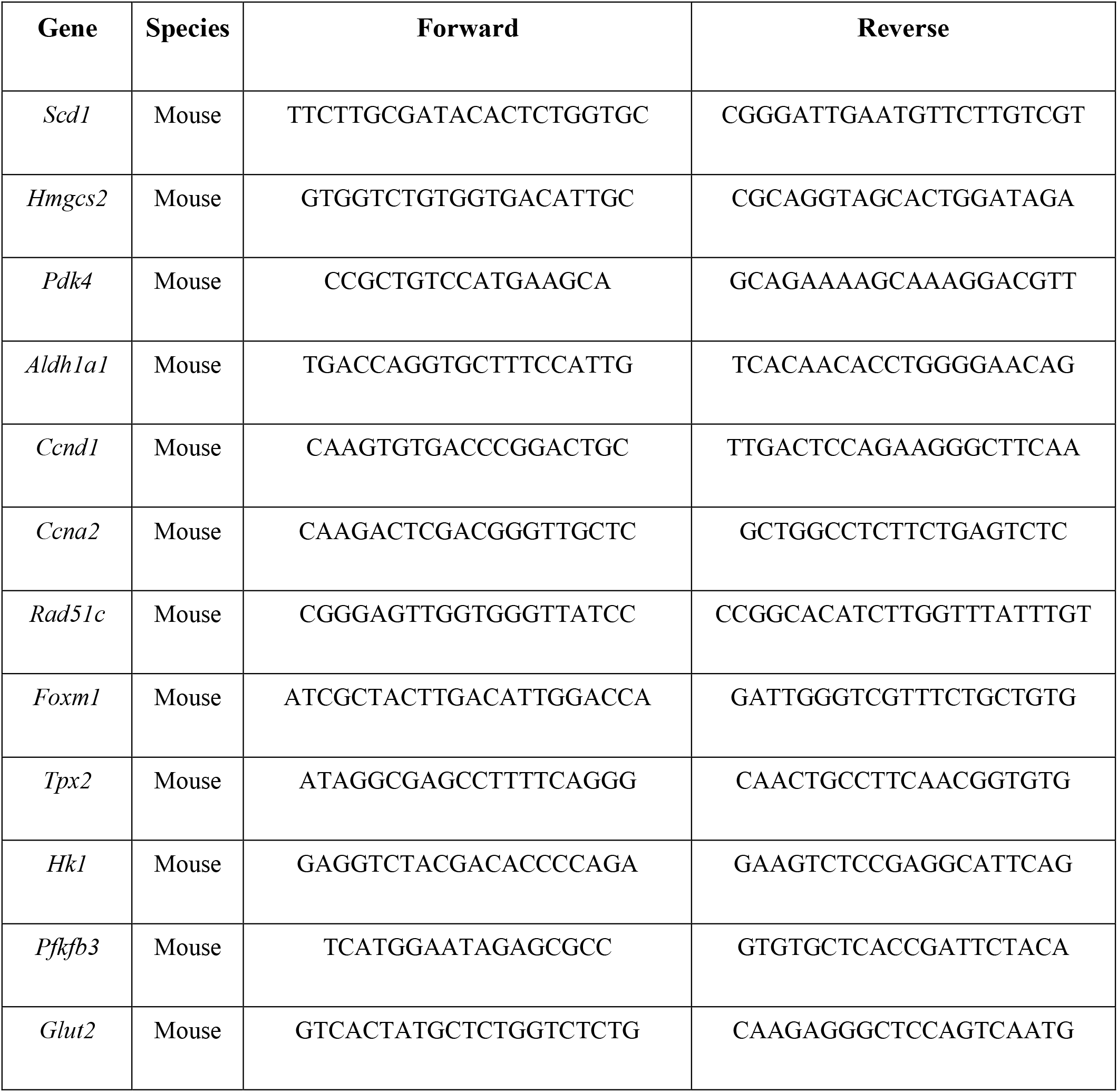
Primers used for qPCR analysis.

## Notes

### Competing Interest Statement

The authors have declared no competing interest.

